# Characterization and inhibitor sensitivity of ARAF, BRAF, and CRAF complexes

**DOI:** 10.1101/2025.08.14.670349

**Authors:** Emre Tkacik, Dong Man Jang, Kayla Boxer, Byung Hak Ha, Michael J. Eck

## Abstract

The RAS-RAF-MEK-ERK signaling pathway controls cellular growth and proliferation, and mutational activation of this pathway is a frequent cause of cancer. Most prominently, the V600E mutation in BRAF causes malignant melanoma, papillary thyroid cancer and other malignancies. Rare but recurrent activating mutations in the other two RAF isoforms, ARAF and CRAF, have also been identified in diverse cancers. Distinct classes of RAF inhibitors have been developed, particularly for BRAF^V600E^, but their potencies against the three RAF isoforms have not been systematically compared. Here we biochemically characterize monomeric and dimeric preparations of ARAF, BRAF, and CRAF and measure the potencies of a panel of thirteen type I, type I.5, and type II RAF inhibitors against each active RAF preparation. Type I inhibitor SB590885 is roughly equipotent across RAF isoforms and, as expected, type I.5 inhibitors are typically most potent against BRAF^V600E^. Despite their reputation as pan-RAF inhibitors, type II inhibitors as a class are potent inhibitors of CRAF but exhibit relative sparing of ARAF and intermediate potencies against BRAF. Type II compounds inhibit BRAF and CRAF with marked positive cooperativity, and their apparent potencies are insensitive to ATP-concentrations. Crystal structures of CRAF in complex with type I.5 inhibitor PLX4720 reveal an asymmetric CRAF dimer with one CRAF subunit bound in the inactive state and the second bound in an αC-helix-in, active conformation with an altered inhibitor pose. Our findings have important implications for understanding the pharmacology of current RAF inhibitors and will inform development of new agents with distinct isoform selectivity.

## Introduction

RAF family kinases (ARAF, BRAF, and CRAF) are the first node in the MAP kinase (MAPK) signaling cascade and are responsible for phosphorylating MEK, which in turn phosphorylates ERK (there are two MEK isoforms MEK1 and MEK2 and also two ERK isoforms ERK1 and ERK2, for simplicity we refer to them collectively as MEK and ERK) (1, 2). ERK is the terminal node in this cascade and, once activated, it phosphorylates diverse transcription factors to regulate cell growth, survival, and proliferation (3). RAF is maintained in an inactive state in the cytoplasm as a complex with its substrate MEK and 14-3-3, a dimeric protein that binds two phosphoserine motifs in the RAF monomer (4–9). RAFs are activated by RAS (KRAS, NRAS or HRAS) (10–14). Active, GTP-bound RAS recruits the RAF:MEK:14-3-3 complex to the plasma membrane, resulting in the formation of active RAF dimers.

Over the past several years, structural studies have revealed the structure of BRAF:MEK:14-3-3 complexes in the autoinhibited and active states (15). In the autoinhibited state, the 14-3-3 dimer binds two phosphoserine sites that flank the kinase domain, precluding its dimerization (16, 17). The N-terminal Ras-binding domain (RBD) and cysteine-rich domains (CRD) also stabilize this state. In the active state, the 14-3-3 dimer rearranges to bind the phosphoserine site in the C-terminus of two RAFs, promoting and stabilizing a back-to-back interaction of their kinase domains (16–19). The RAF and MEK kinase domains interact in a face-to-face orientation, allowing phosphorylation of MEK upon RAF activation (8, 16). Once phosphorylated, MEK dissociates from the active RAF dimer (20).

Somatic mutations in BRAF are a relatively common cause of cancer. The V600E point mutation in BRAF drives more than 50% of malignant melanoma and is also found in thyroid, lung and many other cancers (21, 22). BRAF^V600E^ is active as a monomer and therefore does not depend on RAS-driven membrane recruitment and dimerization for its activation (23). Oncogenic mutations in ARAF and CRAF also occur in cancer, though far less frequently than in BRAF (24, 25). The most common ARAF and CRAF mutations disrupt the negative-regulatory 14-3-3 binding site, thereby precluding the fully autoinhibited state of the kinase (26). Beyond cancer, germline mutations in RAF are found in RASopathies, a group of related developmental syndromes caused by diverse alterations in the MAPK pathway (27). Mutations in CRAF are a cause of Noonan syndrome, while BRAF mutations have been identified in cardiofaciocutaneous syndrome (28, 29).

Many structurally diverse inhibitors have been developed to target aberrantly activated RAFs in cancer (30). Like other ATP-competitive kinase inhibitors, these fall into three classes defined by their binding modes (30, 31). Type I inhibitors bind the active, “αC-helix-in” conformation of the kinase, while type I.5 and type II bind distinct inactive conformations; type I.5 inhibitors bind an “αC-helix-out” conformation while type II inhibitors bind the αC-helix-in, “DFG-out” conformation (30, 31). The αC-helix lies in the N-lobe of the kinase domain and plays a key role in the regulation of many kinases where it adopts an outward position in the inactive state and is rearranged to inward position upon activation, typically as a result of dimerization or activation-loop phosphorylation. The DFG-motif is a three-residue segment (Asp-Phe-Gly) located at the N-terminus of the activation loop and type II inhibitors induce or recognize a “flipped” or “DFG-out” conformation of this segment in which the phenylalanine residue rather than the adjacent aspartic acid extends toward the ATP pocket.

Vemurafenib, dabrafenib and encorafenib are type I.5 inhibitors used to treat BRAF^V600E^-driven malignant melanoma and certain other cancers (32). The type I.5 binding mode of these compounds allows for mutant-selective inhibition of BRAF^V600E^ because this conformation is uniquely accessible when BRAF is in the monomeric state. Accordingly, these molecules are frequently referred to as RAF monomer inhibitors. A disadvantage of this binding mode is that alterations that promote dimerization of BRAF^V600E^, including upstream activation of RAS or splice variants in BRAF^V600E^, confer resistance to these agents (30). RAF dimer inhibitors have also been developed, such as type I inhibitors GDC-0879 and SB590885 and type II inhibitors belvarafenib, naporafenib and tovorafenib. Tovorafenib has recently received approval for treatment of pediatric low-grade gliomas driven by KIAA1549:BRAF, a constitutively dimerized form of BRAF created by its truncation and fusion with a portion of the KIAA1549 open reading frame (33–35).

RAF inhibitors have long been observed to induce activation of wild-type RAF, leading to downstream activation of ERK, a phenomenon referred to as paradoxical ERK activation (or more simply, paradoxical activation) (36). Inhibitors with this property increase ERK activation at lower concentrations but act as inhibitors at higher concentrations, resulting in a bell-shaped dose-response profile. Paradoxical activation is a consequence of inhibitor-induced dimerization (or heterodimerization) of wild-type RAFs (37, 38). This effect is particularly acute with conventional type I.5 inhibitors, as they are ineffective at inhibiting the active RAF dimers they induce (30). In patients, paradoxical activation can lead to development of secondary skin lesions, including squamous cell carcinomas and keratoacanthomas (39, 40). For this reason, these RAF inhibitors are now used in combination with a MEK inhibitor in treatment of BRAF^V600E^ driven cancers, both to overcome this effect and improve efficacy and tolerability (32, 41). More recently developed “paradox-breaking” inhibitors such as PLX8394 also have a type I.5 binding mode, but overcome paradoxical activation by more strongly locking the kinase in the C-helix-out inactive conformation (42).

We are working to better understand the potency and selectivity of RAF inhibitors across RAF dimers and the BRAF^V600E^ variant, to better inform deployment of existing inhibitors and to facilitate development of more selective agents. In an earlier study, we profiled the inhibitory activity of type II Raf inhibitors tovorafenib and naporafenib across the three RAF isoforms using defined, 14-3-3-stabilized dimers of ARAF, BRAF and CRAF with bound MEK1. Here we extend this study with profiling of a panel of 13 inhibitors with type I, type I.5 and type II binding modes. In addition, we describe kinetic characterization of these 14-3-3-bound RAF dimers as well as preparation and characterization of monomeric RAF:MEK complexes. We find that as a class, type II inhibitors are most potent on CRAF with relative sparing of ARAF. Additionally, these compounds inhibit RAF dimers with marked positive cooperativity and are comparatively insensitive to ATP concentrations, despite their ATP-competitive binding modes. Type I inhibitor SB590885 was potent across all three RAF isoforms and dabrafenib potently inhibited RAF dimers in addition to BRAF^V600E^, which is surprising considering that type I.5 RAF inhibitors are generally thought to spare RAF dimers. Finally, co-crystal structures of a CRAF:MEK complex with PLX4720 provide the first views of CRAF with a type I.5 inhibitor and reveal an asymmetric binding mode. Collectively, our studies provide new understanding of the RAF isoform selectivity of diverse inhibitor classes and have important implications for understanding the molecular mechanisms underlying paradoxical activation.

## Results and Discussion

### Characterization of dimeric and monomeric ARAF, BRAF, and CRAF complexes

We previously described preparation of active, dimeric complexes of ARAF, BRAF and CRAF with MEK1 and a 14-3-3 dimer (43). In these constructs, the N-terminal regulatory regions of RAF are truncated, but the C-terminal 14-3-3 binding site is retained, allowing isolation of the active 14-3-3-bound dimer. These BRAF and CRAF constructs co-purified with an endogenous 14-3-3 dimer when expressed using the baculovirus/insect cell system, but efficient production of ARAF required preparation of a chimeric construct in which we fused 14-3-3*ε* to the C-terminus of ARAF (43). For ARAF and CRAF, we prepared constructs in which the N-terminal acidic (NtA) motif, a site of phosphorylation, was mutated to match the SSDD sequence of BRAF, a modification known to increase activity of CRAF (44–47). We co-expressed these constructs in insect cells with a non-activatable variant of MEK1 in which the activation loop serine residues are mutated to alanine (S218A/S222A). The resulting purified complexes are illustrated schematically in Figure 1A. Consistent with our earlier studies of these complexes, all were catalytically active, though the ARAF^SSDD^ and CRAF^WT^ complexes were markedly less active than the BRAF and CRAF^SSDD^ complexes (Figure 1B).

**Figure 1.**
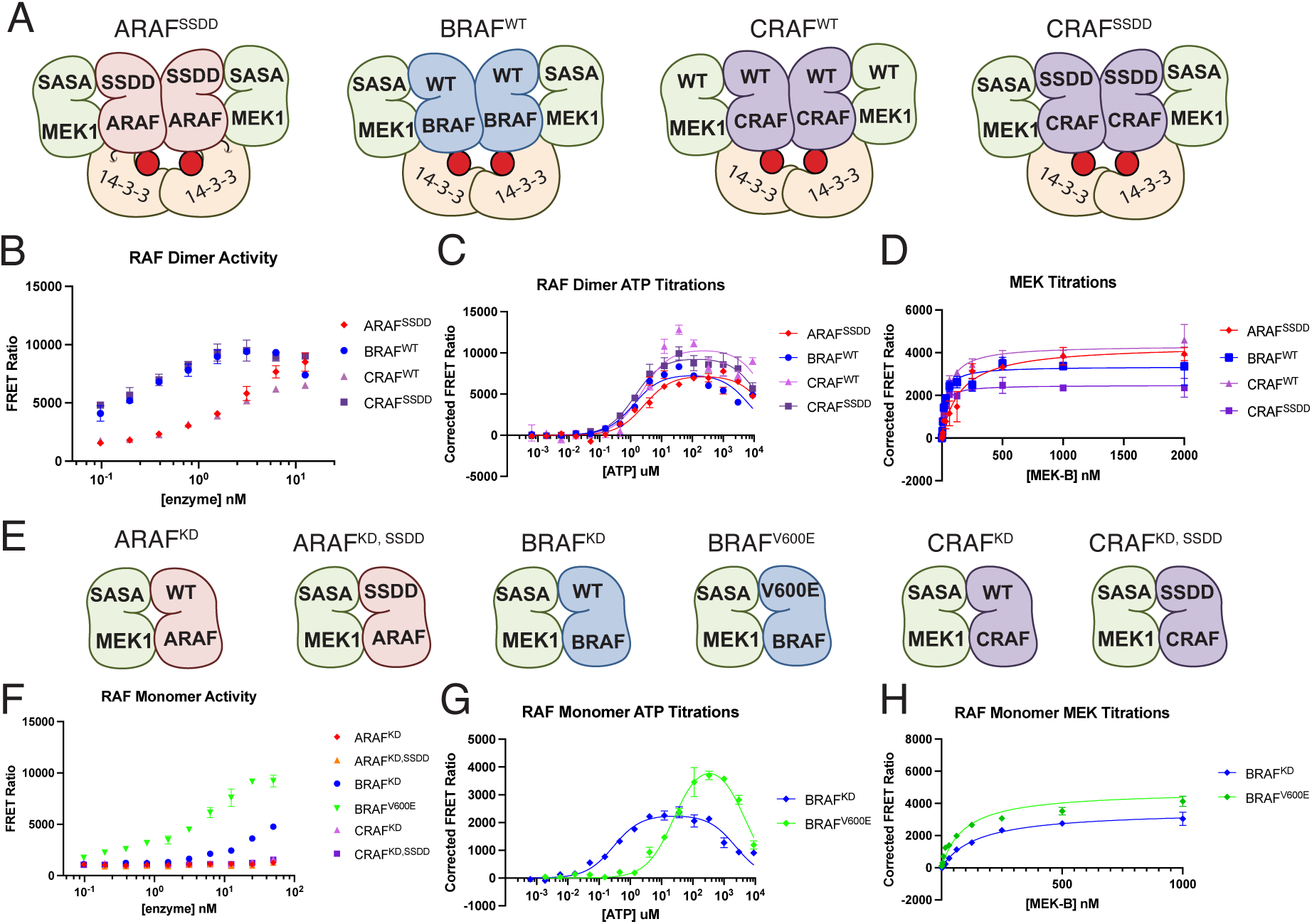
RAF protein constructs and their activity. *A*, schematic of RAF dimer complexes studied here. We refer to these preparations as ARAF^SSDD^, BRAF^WT^, CRAF^WT^, and CRAF^SSDD^, as indicated. *B*, activity of purified RAF dimer complexes as assayed by TR-FRET. Ratio of emission at 665/620 nm is plotted for increasing concentrations of each RAF dimer complex. *C*, Representative ATP titrations for each RAF dimer complex (representative experiment from n≥3). *D*, Representative Michaelis-Menten plots for each RAF dimer complex from MEK titrations (representative experiment from n≥3). *E*, schematic of RAF monomer complexes studied here. We refer to these preparations as ARAF^KD^, ARAF^KD,SSDD^, BRAF^KD^, BRAF^V600E^, CRAF^KD^, and CRAF^KD,SSDD^, as indicated. *F*, activity of purified RAF monomer complexes by TR-FRET. *G*, Representative ATP titrations for BRAF^KD^ and BRAF^V600E^ monomer complexes (representative experiment from n≥3). *H*, Representative Michaelis-Menten plots for BRAF^KD^ and BRAF^V600E^ monomer complexes from MEK titration (representative experiment from n≥3).

In K_m[ATP]_ determinations, we observed marked substrate inhibition with all the RAF proteins tested and accordingly we fit these curves with a substrate inhibition model (Figure 1C,G). The dimeric RAF complexes exhibited closely similar Michaelis constants for ATP, with K_m[ATP]_ values near 1 μM (Table 1). For the inhibitory phase of the titrations, the apparent Ki values were in the millimolar range and were more variable (Table 1). This non-Michaelis Menten behavior has been previously reported for BRAF (19). Unlike typical substrate inhibition that occurs via a second binding site, substrate inhibition by ATP with RAFs likely occurs via binding at the canonical ATP-site of the kinase. The inactive monomeric state of RAFs is stabilized by binding of ATP (16), and ATP-dependent inhibition is overcome by 14-3-3-driven dimerization (19). Higher ATP concentrations (above 50-100 μM) can begin to overcome the activating effect of 14-3-3-driven dimerization. We speculate that high concentrations of ATP promote the inhibited conformation of the kinase domain and may weaken or break the back-to-back dimer interaction of the kinase domains. This could potentially occur without release of the 14-3-3 interaction with the RAF C-terminal tails.

**Table 1.** Enzymatic K_M[MEK]_, K_M[ATP]_ and K_i[ATP]_ values for ARAF, BRAF, and CRAF proteins.*

|  | ARAF<sup>SSDD</sup> | BRAF<sup>WT</sup> | CRAF<sup>WT</sup> | CRAF<sup>SSDD</sup> | BRAF<sup>KD</sup> | BRAF<sup>V600E</sup> |
| --- | --- | --- | --- | --- | --- | --- |
| **$K_{m-MEK}$ (nM)** | 120 ± 50 | 80 ± 40 | 50 ± 10 | 30 ± 10 | 150 ± 10 | 100 ± 30 |
| **$K_{m-ATP}$ (μM)** | 1.58 ± 0.19 | 1.90 ± 0.84 | 1.56 ± 1.50 | 0.73 ± 0.38 | 0.18 ± 0.04 | 12.6 ± 7.80 |
| **$K_{i-ATP}$ (mM)** | >15 | 4.84 ± 2.85 | >15 | >15 | 3.53 ± 0.58 | 6.34 ± 4.07 |

\*Results are reported as mean ± SD ( $n \geq 3$ in triplicate).

### Potency and selectivity of diverse RAF inhibitors across RAF isoforms

To explore the isoform selectivity of different classes of RAF kinase inhibitor, we measured the inhibition of the dimeric 14-3-3-bound RAF complexes described above against a panel of type I, I.5, and II RAF inhibitors. BRAF^WT^ and CRAF^SSDD^ were assayed at a final concentration of 1 nM, while CRAF^WT^ and ARAF^SSDD^ were assayed at 4 and 10 nM respectively. We also tested the monomeric BRAF^V600E^ complex at 1 nM. Substrate MEK1 was present at a concentration of 250 nM and all reactions were preincubated with inhibitor for 40 min, allowed to react at a concentration of 200 μM ATP for 30 min before readout using our TR-FRET assay. The IC_50_ and Hill slope (nH) values of each inhibitor for each enzyme are reported in Table 2 and corresponding inhibition curves are presented in Supplementary Figure 1. Inhibition curves for selected inhibitors are presented in Figure 2.

**Figure 2.**
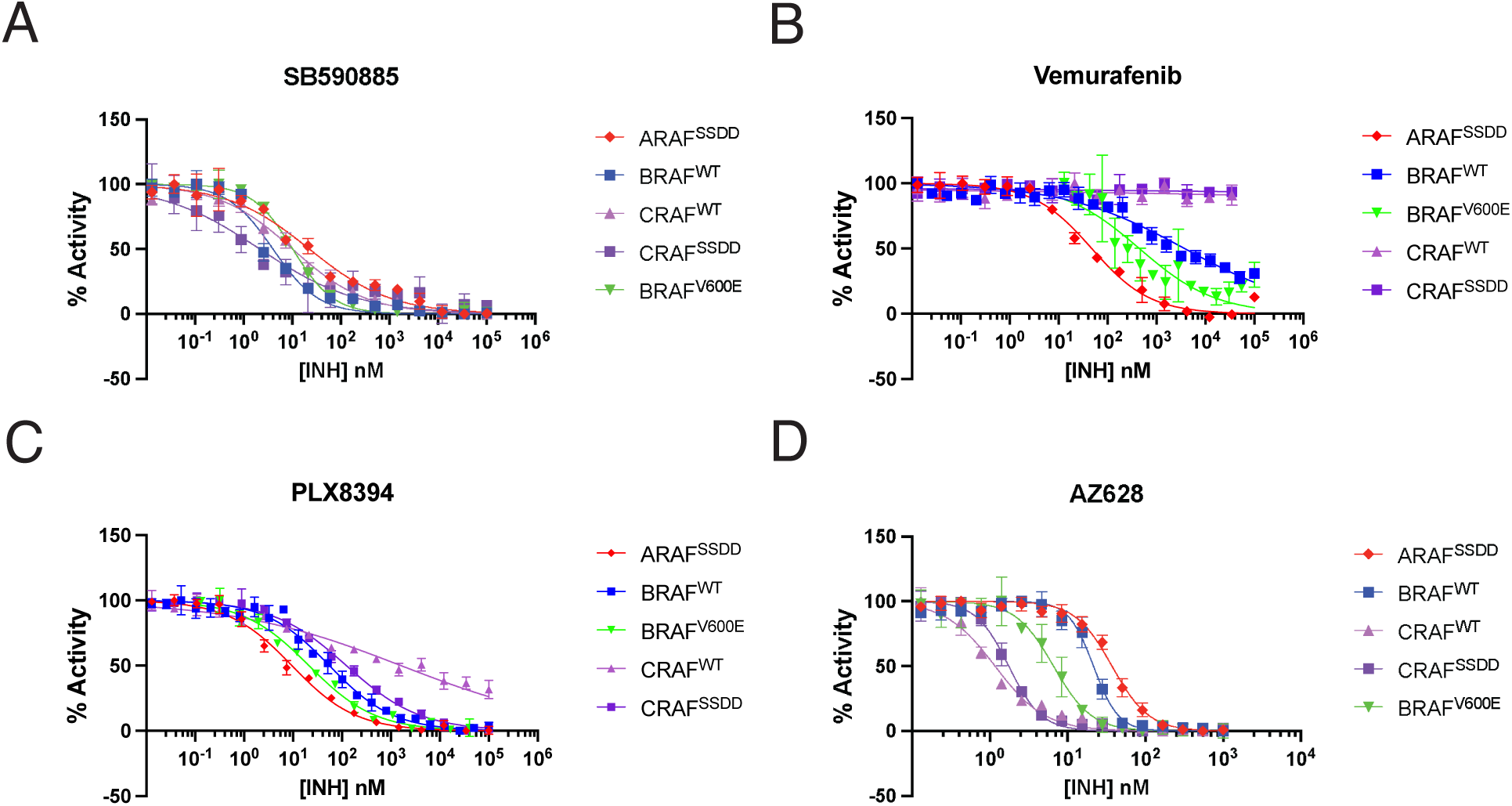
Profiling of representative RAF inhibitors across RAF dimers and BRAF^V600E^. Concentration-response curves for type I inhibitor SB590885 (A), type I.5 inhibitor vemurafenib (B), paradox-breaking type I.5 inhibitor PLX8394 (C), and type II inhibitor AZ628 (D). Data are plotted as mean ± SD from one independent experiment performed in triplicate (n≥3).

**Table 2.** Enzymatic IC_50_ and Hill slope (nH) values for ARAF, BRAF, and CRAF variants.*

|  |  | ARAF<sup>SSDD</sup> |  | BRAF<sup>WT</sup> |  | BRAF<sup>V600E</sup> | CRAF<sup>WT</sup> |  | CRAF<sup>SSDD</sup> |  |
| --- | --- | --- | --- | --- | --- | --- | --- | --- | --- | --- |
|  |  | IC<sub>50</sub><br>(nM) | nH | IC<sub>50</sub><br>(nM) | nH | IC<sub>50</sub><br>(nM) | IC<sub>50</sub><br>(nM) | nH | IC<sub>50</sub><br>(nM) | nH |
| Type I | GDC0879 | 642 ± 61.0 | -0.4 | 291 ± 318 | -0.9 | 553 ± 46.1 | 130 ± 152 | -1.0 | >999 | -0.7 |
|  | SB590885 | 14.6 ± 3.90 | -0.6 | 3.44 ± 0.12 | -1.0 | 10.2 ± 1.04 | 11.6 ± 1.19 | -0.8 | 2.50 ± 1.63 | -0.8 |
| Type I.5 | Vemurafenib | 47.6 ± 9.12 | -0.9 | >999 | -0.6 | 111 ± 33.9 | >999 | -0.7 | >999 | -0.8 |
|  | Dabrafenib | 8.58 ± 5.15 | -1.1 | 9.01 ± 7.66 | -1.4 | 3.26 ± 0.45 | 18.9 ± 24.0 | -0.9 | 23.1 ± 19.0 | -1.0 |
| Type I.5<br>PB | PLX7904 | 899 ± 516 | -0.6 | >999 | -0.6 | 320 ± 82.4 | >999 | -0.4 | >999 | -0.4 |
|  | PLX8394 | 59.6 ± 54.8 | -0.8 | 54.0 ± 40.1 | -0.9 | 21.1 ± 1.78 | >999 | -0.5 | 279 ± 202 | -0.5 |
| Type II | Tovorafenib | 225 ± 67.3 | -1.4 | 208 ± 68.2 | -2.5 | 326 ± 197 | 60.7 ± 37.3 | -1.2 | 57.8 ± 40.0 | -1.8 |
|  | Naporafenib | 161 ± 19.2 | -3.2 | 37.1 ± 23.0 | -2.3 | 51.3 ± 1.16 | 5.46 ± 2.67 | -1.9 | 6.05 ± 2.41 | -2.6 |
|  | Sorafenib | >999 | -0.9 | >999 | -1.2 | >999 | 136 ± 30.8 | -1.2 | 201 ± 61.9 | -1.3 |
|  | Ponatinib | 334 ± 77.0 | -1.2 | 115 ± 32.4 | -2.0 | 21.3 ± 3.58 | 14.9 ± 5.84 | -0.9 | 19.7 ± 6.14 | -1.6 |
|  | AZ628 | 44.1 ± 21.2 | -1.8 | 22.9 ± 4.44 | -3.1 | 7.23 ± 1.92 | 2.05 ± 0.51 | -1.1 | 2.18 ± 0.35 | -2.6 |
|  | Belvarafenib | 69.9 ± 32.0 | -1.3 | 19.9 ± 0.98 | -3.0 | 13.3 ± 0.87 | 5.05 ± 1.81 | -1.6 | 5.66 ± 1.12 | -3.0 |
|  | LY3009120 | 52.0 ± 45.5 | -3.6 | 17.1 ± 3.86 | -2.2 | 7.75 ± 2.14 | 3.10 ± 0.94 | -1.4 | 3.07 ± 0.38 | -2.9 |

\* BRAF<sup>V600E</sup> was fit with a 3-parameter model, so no Hill slopes are shown. Experiments were performed in triplicate in at least three independent experiments (n≥3).

Between the two type I inhibitors tested, SB590885 was by far the more potent, with IC_50_ values in the range of 2.5 nM against CRAF^SSDD^ to ∼15 nM against ARAF (Figure 2A, Table 2). Of all inhibitors tested, it was also the most equipotent across RAF isoforms. While GDC0879 inhibited BRAF^WT^ and CRAF^WT^ with IC_50_ values in the range of 100 – 300 nM, it exhibited IC_50_ values of greater than 500 nM for ARAF^SSDD^ and BRAF^V600E^ and more than 1 μM against CRAF^SSDD^ (Table 2).

We tested a total of four type I.5 inhibitors, including two (PLX7904 and PLX8394) that are referred to as “paradox breaking” inhibitors because they do not induce paradoxical ERK activation in cells (42). All four type I.5 agents were potent against the monomeric BRAF^V600E^ enzyme. This is as expected, because the C-helix-out binding mode of type I.5 inhibitors is antagonized by dimerization, which activates RAF by promoting the C-helix-in conformation. Vemurafenib was active against BRAF^V600E^ (∼111 nM) but spared WT BRAF and CRAF dimers (IC_50_ > 1 μM). Unexpectedly, vemurafenib was most potent against ARAF dimers (∼48 nM). Dabrafenib was considerably more potent that vemurafenib against BRAF^V600E^ (3 nM), and considering its type I.5 binding mode, it exhibited surprising activity against RAF dimers (9 nM, 9 nM, and 19 nM for ARAF, BRAF, and CRAF, respectively). Dabrafenib’s potency against these dimers may be explained, at least in part, by a crystal structure with dabrafenib bound to BRAF that shows its binding is compatible with dimerization with modest rearrangements of the dimer interface (Supplementary Figure 2). Paradox breaker PLX8394 inhibited BRAF^V600E^ with an IC_50_ of ∼20 nM and was about 3-fold less active on BRAF^WT^ and ARAF^SSDD^ dimers. Interestingly, it was not potent against CRAF^WT^ dimers (IC_50_ > 1 μM). PLX7904 was most potent against BRAF^V600E^, with an IC_50_ of 320 nM and IC_50_ values in the micromolar range against other RAFs.

We profiled seven type II RAF dimer inhibitors (Supplemental Figure 1, Figure 2D), all of which were most potent against CRAF and relatively sparing of ARAF (Table 2). Naporafenib, AZ628, belvarafenib, and LY3009120 all exhibited single-digit nanomolar potency against CRAF^WT^ and CRAF^SSDD^. The ARAF-sparing nature of naporafenib, tovorafenib, and belvarafenib has been previously noted (26, 43, 48, 49), and we find here that this property pertains to all the structurally diverse type II inhibitors studied here. All type II compounds tested displayed at least 10-fold greater potency against CRAF dimers than ARAF dimers, except tovorafenib, which was ∼4-fold more potent on CRAF than ARAF. This property was most pronounced for naporafenib, which is approximately 30-fold more potent on CRAF than ARAF. Considering the requirement for a DFG-out conformation for binding of type II compounds and the commonality of ARAF-sparing across structurally diverse inhibitors, we expect that their diminished potency on ARAF stems from a higher energetic barrier to adopting this conformation in ARAF as compared with BRAF and CRAF.

Another common feature of the type II inhibitors is that they inhibit BRAF and CRAF^SSDD^ dimers with clear positive cooperativity. This is evident in the steep slope of the dose-response curves and the values of Hill constants obtained with four-parameter fitting of these curves (Figure 2D and Table 2, Hill constants less than -1.0 indicate positive cooperativity). As expected, there was no obvious cooperativity with monomeric BRAF^V600E^ and accordingly three-parameter fits were applied for these inhibition curves. With ARAF^SSDD^ only naporafenib, AZ628 and LY3009120 exhibited clear positive cooperativity. Interestingly, positive cooperativity was markedly diminished or entirely absent for these inhibitors on CRAF^WT^ dimers, despite striking concordance of their potency on CRAF^WT^ versus CRAF^SSDD^ (Table 2). We speculate that the SSDD substitution may lower the energy barrier to conformational rearrangements that accompany dimerization and cooperative binding of type II inhibitors.

### BRAF inhibition by type II inhibitors is insensitive to ATP concentration

We conducted additional inhibitor titrations of BRAF^WT^ dimers with selected compounds at ATP concentrations of 10 *μ*M, 100 *μ*M, and 1000 *μ*M (Figure 3). The effect of ATP concentration on the IC_50_ of an ATP-competitive inhibitor can be predicted using the Cheng-Prusoff equation: *IC*_50_= *K_i_* × (1 + ([*S*]⁄*K_m_*)) where [S] is the ATP concentration used for an experiment, K_i_ is a given inhibitor’s inhibition constant, and K_m_ is the enzyme’s ATP affinity. Based on the K_m, ATP_ of BRAF^WT^, it is reasonable to expect a 5-10-fold increase in IC_50_ for every 10-fold increase in ATP concentration. The IC_50_ values of type I inhibitor SB590885 and type I.5 inhibitor PLX8394 shifted as expected with increasing concentrations of ATP (Table 3).

**Figure 3.**
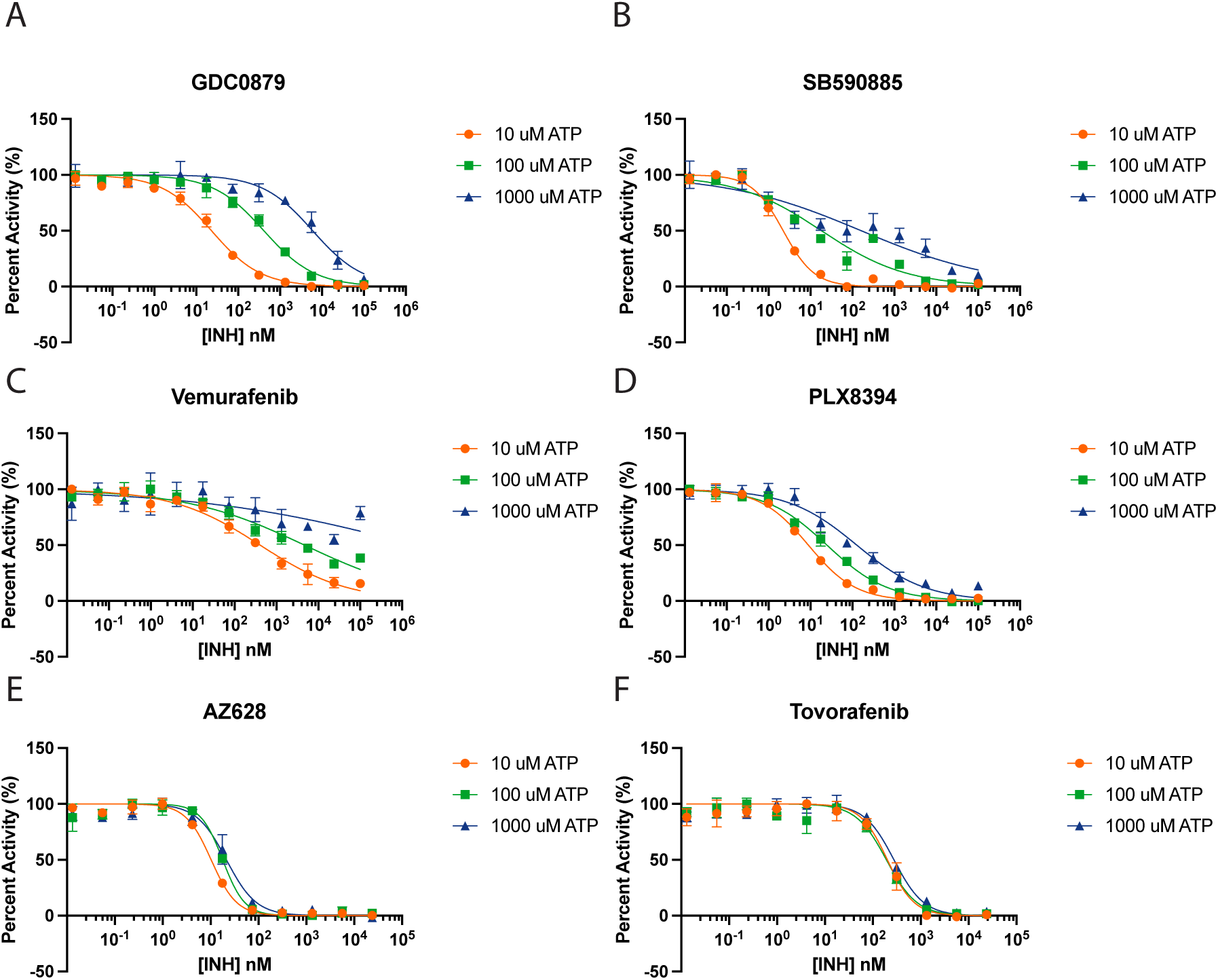
Effect of ATP concentration on inhibition of BRAF^WT^ dimers by selected RAF inhibitors. Inhibition of wild-type BRAF dimers by the indicated inhibitors was measured at ATP concentrations of 10 μM, 100 μM, or 1000 μM in the TR-FRET assay. A, GDC0879; B, SB590885; C, vemurafenib; D, PLX8394; E, AZ628; and F, tovorafenib.

**Table 3.** Enzymatic IC_50_ values of BRAF^WT^ at 10 μM, 100 μM, and 1000 μM ATP.*

| IC<sub>50</sub> (nM) | 10 μM ATP | 100 μM ATP | 1000 μM ATP |
| --- | --- | --- | --- |
| **GDC0879** | 21.7 ± 3.96 | 276 ± 171 | >3000 |
| **SB590885** | 1.66 ± 0.43 | 6.68 ± 5.00 | 57.5 ± 66.6 |
| **Vemurafenib** | 249 ± 173 | >2000 | >10000 |
| **Dabrafenib** | 2.71 ± 0.72 | 5.58 ± 1.55 | 13.3 ± 5.41 |
| **PLX7904** | 170 ± 121 | >1000 | >10000 |
| **PLX8394** | 11.1 ± 7.84 | 38.5 ± 34.2 | 259 ± 316 |
| **Tovorafenib** | 140 ± 64.8 | 158 ± 36.6 | 280 ± 56.0 |
| **Naporafenib** | 29.1 ± 8.31 | 38.2 ± 11.4 | 54.9 ± 16.5 |
| **AZ628** | 16.6 ± 11.9 | 35.7 ± 33.7 | 38.9 ± 30.4 |
| **Belvarafenib** | 14.2 ± 6.36 | 21.5 ± 9.23 | 29.3 ± 8.75 |

\*Inhibitor titrations were performed in triplicate in three independent experiments (n ≥ 3).

By contrast, the IC_50_ values of type II inhibitors tovorafenib, naporafenib, AZ628, and belvarafenib were only modestly affected by increasing ATP concentrations. Type II inhibitors are clearly ATP-competitive, in that they occupy the ATP binding cleft as seen in numerous co-crystal structures (43, 48). Also, the DFG-out conformation induced by naporafenib and other type II inhibitors is not compatible with binding of ATP. Thus we suspect that the apparent non-competitive behavior observed here arises from the type II binding mode and likely also the cooperativity of inhibitor binding to the BRAF dimer. To our knowledge, this behavior has not been previously reported for type II RAF inhibitors.

### Crystal structure of CRAF in complex with PLX4720

While there are more than 100 structures of BRAF available in the Protein Data Bank (PDB), there is scant structural information available for CRAF. Recent cryo-EM structures provided views of CRAF in its apo form and bound to type I inhibitor GDC-0879 (50, 51), but to our knowledge no structures are available with type I.5 or type II inhibitors. Furthermore, our examination of an earlier crystal structure with type I inhibitor GDC0879 (PDB ID: 3OMV) (38) reveals that it is actually a BRAF structure that was inadvertently interpreted and modeled as CRAF (Supplementary Figure 3).

We co-crystallized the CRAF:MEK complex with PLX4720, a type I.5 inhibitor that is a close analog of vemurafenib. Crystallization was facilitated by inclusion of CH5126766, a MEK inhibitor that is known to stabilize RAF:MEK complexes (52). The complex crystallized in space group *P*6_5_22 with two essentially identical copies of a CRAF_2_MEK_2_ heterotetramer in the asymmetric unit (RMSD = 0.715 Å). We refined the structure to an R-value of 23.02% (R_free_ = 0.2639) using data extending to a resolution of 2.91 Å (Supplementary Table 1). The structure reveals clear electron density for both PLX4720 and CH5126766 (Supplementary Figure 4A-C).

In this structure, CRAF forms a back-to-back dimer, with MEK1 bound to each CRAF kinase domain in a face-to-face orientation as previously seen in active-state BRAF and CRAF complexes with MEK1 (Figure 4A). Unlike prior structures of CRAF_2_MEK_2_ heterotetramers (50, 51), only one CRAF protomer adopts an active, αC-helix-in conformation. In the other protomer, PLX4720 forces an inactive, αC-helix-out conformation. In the first protomer, PLX4720 binds in the type 1.5 manner expected based on prior structures in complex with BRAF (Figure 4B). In the opposite protomer, however, it exhibits a non-canonical binding mode in which the pyrrolopyridine scaffold is flipped and the compound adopts a compact C-shaped conformation such that the propylsulfonamide packs below and adjacent to the pyridine ring, forming hydrogen bonds with both the active site lysine (K375) and the main chain amide of D486 in the DFG motif (Figure 4C). This altered pose allows CRAF to adopt the active, αC-in conformation (Figure 4C). In our initial CRAF_2_MEK_2_ crystal structure, the occupancy of the ATP analog in the MEK1 active site was low, resulting in an apparent crystallographic artifact in which the activation loop of CRAF extended into the ATP binding site of MEK1 (Supplementary Figure 4D,E). A subsequent structure of the same complex crystallized with a higher concentration of AMPPNP (2 mM) resolved this artifact and showed clear density for the nucleotide in the MEK1 active site (Supplementary Figure 4D,F).

**Figure 4.**
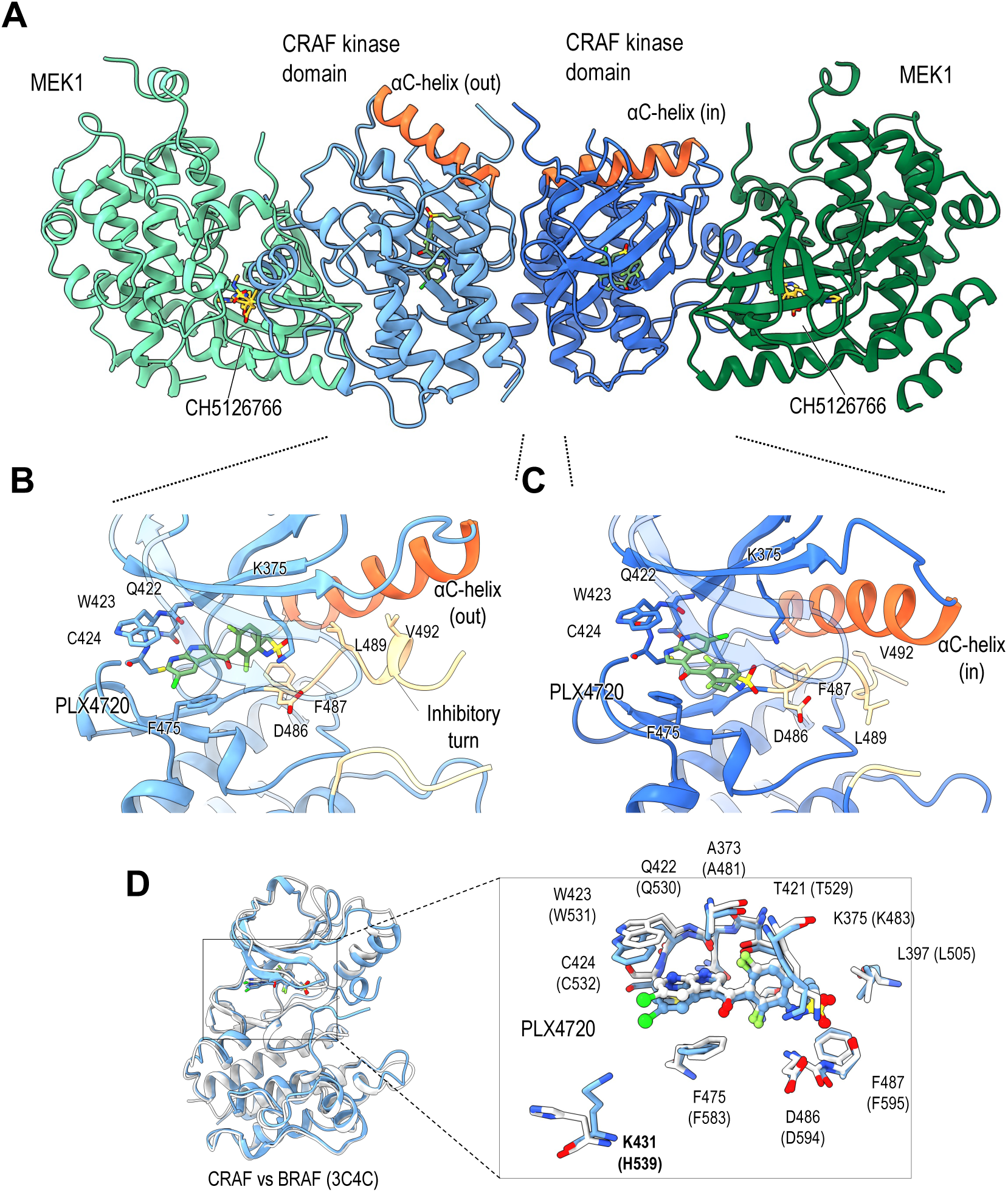
Crystal structure of CRAF with vemurafenib analog PLX4720. A, Ribbon representation of the MEK:CRAF:CRAF:MEK heterotetramer. MEK is colored green, CRAF is colored blue, the CRAF αC-helices are colored orange. MEK inhibitor CH5126766 is bound to each MEK protomer, and is colored yellow. PLX4720 is bound to each CRAF kinase and is colored green. The orange αC-helices of each CRAF protomer are in different conformations. B, Detailed view of PLX4720 bound to CRAF in the αC-out conformation. The αC-helix is colored orange, the inhibitory turn is colored tan, and the inhibitor is shown as sticks in green. C, Detailed view of PLX4720 bound to CRAF in the αC-in conformation. The αC-helix is colored orange, no inhibitory turn is present due to the αC-in conformation, but the relevant residues are colored tan, and the inhibitor is shown as sticks in green. D, PLX4720 bound CRAF in the αC-out conformation overlayed with crystal structure of BRAF bound to PLX4720 (PDB ID: 3C4C) (53). Full kinase domains are shown on the left, with a detailed view in the box on the right. PLX4720 takes similar binding poses in each structure, as do the active site residues. Non-conserved lysine 431 (H539 in BRAF) is highlighted in bold text. CRAF is colored blue, BRAF is colored white.

Comparison of the present structure with that of PLX4720 bound to BRAF (PDB ID: 3C4C) (53) reveals an essentially identical binding mode in the first protomer, with all of the residues contacting the inhibitor conserved between BRAF and CRAF. The one notable difference is K431, which is a histidine in both BRAF and ARAF. Though not in direct contact with the inhibitor, its εN is ∼ 4.5Å from the chlorine substituent on PLX4720 (Figure 4D). Considering recent interest in covalent targeting of lysine residues, this unique feature of CRAF may provide an attractive avenue to development of a CRAF selective agent. Interestingly, this lysine hydrogen bonds with the sidechain of E102 of MEK1, which may increase its nucleophilicity and susceptibility to covalent modification. The BRAF dimer with PLX4720 is also asymmetric, but the inhibitor in the second protomer is accommodated by a flip of the DFG motif and reorientation of the propylsulfonamide substituent of the inhibitor (53).

### Concluding Remarks

To date, RAF inhibitor development has focused primarily on BRAF and mutant-selective inhibitors of BRAF^V600E^ represent a landmark advance in precision medicine (22, 54). The discovery of the unique requirement for CRAF in KRAS-driven tumorigenesis has made CRAF an important target for inhibitor development (55–57), and likewise the role of ARAF in promoting resistance to belvarafenib in melanoma has heightened interest in ARAF as a target (26). While mutant-selectivity is not relevant in these contexts, an understanding of isoform-selectivity is crucial. Because complete blockade of RAF signaling is not tolerable systemically, pharmacologic inhibition should focus on the isoforms involved in signaling in a particular oncogenic context in order to achieve a therapeutic window. Our results show that the ARAF-sparing properties recently reported for selected type II inhibitors are a general property of this class, as is their greater potency against CRAF as compared with BRAF. Thus, the common conception of type II compounds as “pan-RAF” inhibitors is not accurate. Rather, we find that among the inhibitors we evaluated, type I inhibitor SB590885 is most equi-potent across RAF isoforms.

Our results are also relevant to understanding the mechanism of paradoxical activation by RAF inhibitors. Inhibitor binding induces RAF dimerization (58), and a prevailing model for paradoxical activation posits that inhibitor binding to one protomer of a RAF dimer results in an active, but inhibitor resistant conformation in the second protomer (23, 59). This model implies negative cooperativity of inhibitor binding and provides an explanation for why an inhibitor-induced dimer can nevertheless remain active. A crystal structure of BRAF bound to vemurafenib provides support for this model in that it shows the compound bound to one subunit of a RAF dimer in the expected type I.5, αC-helix-out binding mode, while the active site of the opposite protomer remains empty and exhibits an active, αC-helix-in conformation that should not be amenable to inhibitor binding (PDB ID: 3OG7) (60). The present CRAF structure further establishes the plausibility of this model for type I.5 inhibitors by extending it to CRAF. However, our results also call into question the generality of this model. Type II RAF inhibitors are also known to promote RAF dimerization and exhibit paradoxical ERK pathway activation (59), but our results show that they inhibit RAF dimers with marked positive, rather than negative, cooperativity. Mechanistic studies of paradoxical activation by type I and II RAF inhibitors is an area of active investigation in our laboratory and will be the focus of a future manuscript.

## Materials and Methods

### RAF Dimer Expression and Purification

To prepare RAFs as enzymes for assays, SF9 insect cells were co-infected with two baculoviruses, one expressing a MEK construct and the other corresponding to the RAF isoform required for the desired enzymatic complex. MEK1^SASA^:ARAF^SSDD^-14-3-3*ε* was purified using a ARAF^274-606, G300S, Y301/302D^-14-3-3*ε* with n-terminal His_6_ and StrepII tags (ARAF^SSDD^-14-3-3*ε*) construct and a full length His_6_-MEK1^S218/222A^ construct (MEK1^SASA^). Cells were harvested 72 hours post infection and lysed in Ni binding buffer (50 mM Tris pH 7.5, 150 mM NaCl, 5 mM MgCl_2_, 1 mM TCEP, 1 *μ*M AMP-PNP, and protease inhibitor cocktail from Thermofischer Scientific) via sonication. Clarified lysate was removed and bound to equilibrated Ni-NTA agarose beads (Qiagen) by gravity flow, washed with Ni binding buffer supplemented with 30 mM imidazole, then eluted with Ni binding buffer supplemented with 300 mM imidazole. Elutions containing the expressed proteins were pooled and bound to an equilibrated 5 ml StrepTrap HP column (GE Healthcare Life Sciences), washed with Ni binding buffer, then eluted using binding buffer supplemented with 5 mM desthiobiotin. Fractions were analyzed by SDS PAGE and those containing the desired complex were pooled and concentrated to 1 ml by Amicon Ultra concentrator (30 MWCO, Millipore) before being injected onto a Superose 6 10/300 (GE Healthcare Life Sciences) column. SDS PAGE analysis indicates that MEK1^SASA^ and ARAF^SSDD^-14-3-3*ε* co-eluted together along with some endogenous insect 14-3-3, forming a MEK1^SASA^:ARAF^SSDD^-14-3-3*ε* RAF dimer complex that exhibits kinase activity in our TR-FRET assay.

MEK1^SASA^:BRAF^WT^:14-3-3 was prepared using a BRAF^419-766^ construct with an n-terminal His_6_ tag (BRAF^WT^) and the full length MEK1^SASA^ construct described above. Again, approximately 72 hours after infection cell cultures were harvested and lysed in Ni binding buffer (50 mM Tris, 150 mM NaCl, 10 mM MgCl_2_, 1 mM TCEP, 1 *μ*M AMP-PNP, and protease inhibitor cocktail from Thermofischer Scientific) as described above. Clarified lysate was bound to Ni-NTA agarose beads (Qiagen) by gravity flow, washed with Ni binding buffer supplemented with 30 mM imidazole, then eluted with Ni binding buffer supplemented with 250 mM imidazole. Elutions containing expressed proteins were concentrated to 5 ml by Amicon Ultra concentrator (30 MWCO, Millipore) and injected onto a Superdex 200 Increase 10/300 column (GE Healthcare Life Sciences). SDS PAGE analysis of the resulting fractions indicates that the co-expressed MEK1^SASA^ and BRAF^WT^ co-elute with approximately stoichiometric amounts of endogenous insect 14-3-3, to form a highly active MEK1^SASA^:BRAF^WT^:14-3-3 dimer complex, as indicated by TR-FRET. MEK1^SASA^:BRAF^V600E^ monomer complex was obtained by co-infecting insect cells with recombinant baculovirus expressing either full length MEK1^SASA^ or BRAF^445-723, V600E^ (BRAF^V600E^) with an n-terminal His_6_ tag and a c-terminal intein tag. As with the complexes described thus far, cells were harvested approximately 72 hours post infection and sonicated in lysis buffer (25 mM Tris, 100 mM NaCl, 5 mM MgCl_2_, 1 mM TCEP, 1 *μ*M AMP-PNP, and protease inhibitor cocktail from Thermofischer Scientific). After ultracentrifugation at 40,000 rpm, clarified lysate was bound to Ni-NTA agarose beads (Qiagen), washed with lysis buffer supplemented with 25 mM imidazole, then eluted with 500 mM imidazole. Elutions were pooled and incubated with 50 mM MESNA overnight, then the applied to Chitin beads. The flowthrough was concentrated to 1 ml by Amicon Ultra concentrator (10 MWCO, Millipore) then further purified by gel filtration using a Superdex 200 10/300 column (GE Healthcare Life Sciences). SDS PAGE analysis indicated that these constructs eluted together as a MEK1^SASA^:BRAF^V600E^ monomer, without 14-3-3, with high levels of kinase activity.

For MEK1:CRAF^WT^:14-3-3 dimers, insect cell cultures were co-infected with baculoviruses expressing his_6_-MEK1^35-393^ (MEK1) with a c-terminal avi-tag and his_6_-CRAF^308-648^ (CRAF^WT^). This complex was purified like MEK1^SASA^:BRAF^WT^:14-3-3, using identical buffers and purification methods. Briefly, lysates were bound to Ni-NTA beads, washed, and eluted. After SDS PAGE, elutions containing the complex were concentrated to 1 ml and injected onto a Superdex 200 10/300 column for gel filtration. These fractions were run on an SDS PAGE gel, which indicated that MEK1 and CRAF^WT^ co-elute with endogenous 14-3-3. Fractions corresponding to the MEK1:CRAF^WT^:14-3-3 dimer complex were pooled and concentrated for storage at -80 °C until needed for assays. The MEK1^SASA^:CRAF^SSDD^:14-3-3 complex was prepared by co-infecting insect cells with the MEK1^SASA^ expressing baculovirus described above in combination with a his_6_-StrepII-CRAF^308-648, Y340/341D^ (CRAF^SSDD^) expressing baculovirus. As with the other preparations, cell cultures were harvested, lysed, and ultracentrifuged, then clarified lysates were bound, washed, and eluted from Ni-NTA agarose beads using the buffers described above for MEK1^SASA^:BRAF^WT^:14-3-3 purification (50 mM Tris, 150 mM NaCl, 10 mM MgCl_2_, 1 mM TCEP, 1 *μ*M AMP-PNP, and protease inhibitor cocktail from Thermofischer Scientific). Elutions containing both expressed proteins and endogenous 14-3-3 were bound to a prepacked StrepTrap HP column (GE Healthcare Life Sciences), washed briefly, then eluted using binding buffer supplemented with 5 mM desthiobiotin. Elutions containing the complex were pooled and concentrated to 5 ml by Amicon Ultra concentrator (30 MWCO, Millipore) then injected onto a Superdex 200 Increase 10/300 column. This complex eluted with endogenous insect 14-3-3 proteins as an active MEK1^SASA^:CRAF^SSDD^:14-3-3 dimer.

### RAF Monomer Expression and Purification

RAF:MEK monomer complexes were expressed and purified comparably to the active complexes described thus far, but using truncated RAF kinase domain constructs that are incapable of 14-3-3 binding. RAF:MEK monomer complexes were expressed and purified comparably to the active complexes described thus far, but using truncated RAF kinase domain constructs that are incapable of 14-3-3 binding. BRAF^V600E^:MEK1^SASA^ was purified from insect cells coinfected with baculoviruses expressing full-length MEK1^SASA^ and his_6_-BRAF^445-723, V600E^-intein. Cells were centrifuged three days after infection, lysed by sonication using the lysis buffer stated above (50 mM Tris, 150 mM NaCl, 10 mM MgCl_2_, 1 mM TCEP, 1 *μ*M AMP-PNP, and protease inhibitor cocktail from Thermofischer Scientific), then ultracentrifuged for 1-2 hours at 40,000 rpm. The clear lysate was removed from ultracentrifuge tubes and bound by gravity flow to equilibrated Qiagen Ni-NTA agarose beads, washed with 25 mM imidazole, and finally eluted with 500 mM imidazole. Ni-NTA fractions were analyzed by SDS page, and appropriate fractions were pooled and incubated with MESNA overnight at a final concentration of 50 mM. The following morning, this volume was applied to Chitin beads, the flowthrough was collected, then concentrated to 1 ml by Amicon Ultra concentrator (10 MWCO, Millipore). The final purification step was gel filtration on a Superdex 200 10/300 (GE Healthcare Life Sciences), after which fractions were again analyzed by SDS-page. The BRAF^V600E^:MEK1^SASA^ complex eluted together as a highly active heterodimer, as determined by TR-FRET.

For BRAF^KD^:MEK1^SASA^ monomers, insect cells were co-infected with his_6_-BRAF^445-723^-intein (BRAF^KD^) and the full length MEK1^SASA^ construct used frequently in preparation of the RAF dimers. Cell cultures were harvested by centrifugation after 85 hours, resuspended in lysis buffer, sonicated, and ultracentrifuged at 40,000 rpm. The supernatant was bound to Ni-NTA beads, washed, and eluted, then incubated in 50 mM MESNA overnight, to ensure complete self-cleavage of the intein tag. The next day, this volume was concentrated to 5 ml by Amicon Ultra concentrator (30 MWCO, Millipore) then injected onto a Superdex 200 10/300 column, eluting as a BRAF^KD^:MEK1^SASA^ monomer.

ARAF^KD^:MEK1^SASA^ monomers were also purified from insect cells co-infected with recombinant baculoviruses expressing full length MEK1^SASA^ proteins and his_6_-ARAF^282-579^ (ARAF^KD^). After harvesting, cells were lysed and ultracentrifuged as usual, then the clarified lysate was bound to Ni-NTA beads, washed with 30 mM imidazole, and eluted with 300 mM imidazole. Fraction containing the desired complex were pooled and concentrated to 1 ml by Amicon Ultra concentrator (30 MWCO, Millipore). This was injected onto a Superdex 200 10/300 column and eluted as an ARAF^KD^:MEK1^SASA^ a monomer with no basal activity, as evaluated by TR-FRET. CRAF^KD^:MEK1^SASA^ monomers were prepared similarly, by co-expressing full-length his_6_-MEK1^SASA^ with StrepII-CRAF^324-618^. This complex, like others, was first bound to Ni-NTA beads. After washing, elutions with the complex were pooled and diluted with strep binding buffer to a final concentration below 200 mM imidazole. This volume was bound to Streptactin Macroprep beads at 1 ml/min, washed briefly (less than 10 ml) and eluted with 5 mM desthiobiotin. Strep elutions were concentrated to 1 ml, and injected onto an equilibrated Superdex 200 10/300. This purification resulted in CRAF^KD^:MEK1^SASA^ monomers with no basal activity.

ARAF^KD,SSDD^:MEK1^SASA^ and CRAF^KD,SSDD^:MEK1^SASA^ monomers were prepared identically, by co-infecting full-length MEK1^SASA^ with either his_6_-MBP-TEV-ARAF^274-606, G300S, Y301/302D^ (ARAF^KD,SSDD^) or his_6_-MBP-TEV-CRAF^314-618, Y340/341D^ (CRAF^KD,SSDD^). Purification up until N-NTA proceeded as has been described above, at which point Ni elutions were pooled, buffer exchanged, incubated with TEV protease overnight for cleavage of the MBP tag, and passed over Ni-NTA beads a 2^nd^ time the following day. The flowthrough from this was concentrated to 1 ml and injected onto an equilibrated Superdex 200 10/300. Purification in this manner resulted in monomeric ARAF^KD,SSDD^:MEK1^SASA^ and CRAF^KD,SSDD^:MEK1^SASA^ complexes with nearly no activity.

### Kinase Assays

Inhibition assays were performed using a modified HTRF KinEASE tyrosine kinase assay kit (Cisbio). Rather than the provided kit substrate, we purified MEK1^35-393^ and biotinylated it (MEK-B) in house using birA enzyme. Inhibitors were dispensed into black 384-well plates using an HP300e dispenser and normalized to 1% final DMSO concentration per well. Kit assay buffer was supplemented with purified RAF at a final concentration of 1 nM for MEK1^SASA^:BRAF^KD^:14-3-3 and MEK1^SASA^:CRAF^SSDD^:14-3-3, 4 nM for MEK1:CRAF^KD^:14-3-3, 10 nM for MEK1^SASA^:ARAF^SSDD^-14-3-3*ε*, and 25 nM for each of the remaining RAF constructs, as well as purified biotinylated MEK-B at a final concentration of 250 nM. Supplemented kinase buffer was dispensed into 384 well plates using a Multidrop combi dispenser and incubated with inhibitors at room temperature for 40 min before reactions were initiated by addition of 400 μM ATP dispensed using the Multidrop combi dispenser, to a final concentration of 200 μM. Plates were quenched after 30 min at room temperature using the kit detection buffer supplemented with XL665 and an Anti-phospho MEK1/2-Eu antibody (Cisbio). The FRET signal ratio was measured at 665 and 620 nm using a PHERAstar microplate reader and processed using GraphPad Prism fit to a three-parameter dose-response model with Hill Slope constrained to -1 and a four-parameter dose-response model that fits the Hill Slope to the data. Assays were performed in triplicate three independent times.

For enzyme and ATP titrations, the experimental setup followed that above, varying ATP or enzyme concentration as necessary for the corresponding titration. ATP titrations were fit with substrate inhibition curves in GraphPad Prism to determine K_m, ATP_ and K_i, ATP_. Changing the substrate concentration without changing the corresponding detection reagent concentration (XL665) results in artificial decreases in observed FRET at high MEK concentrations. To avoid this effect, MEK titrations were performed at a constant MEK-B:XL665 ratio, rather than a constant [XL665]. In all experiments, the MEK-B:XL665 ratio is 4:1 (250 nM MEK-B and 62.5 nM XL665), in the MEK titrations we maintain this ratio as MEK is increased/decreased, which results in classic Michaelis Menten curves. If XL665 is held constant, and instead fit with substrate inhibition, the resulting K_m, MEK_ values are similar.

### MEK Expression and Purification for Assay Substrate

MEK for use in the TR-FRET assay as substrate was expressed in insect cells with His_6_-MEK1^35-393^ (MEK1) expressing recombinant baculovirus. This construct also contains a c-terminal avi-tag to allow for biotinylation by birA, which was expressed and purified from *E. coli* in house. Cell cultures were pelleted, lysed, then ultracentrifuged at 40,000 rpm in Ni binding buffer (50 mM Tris, 150 mM NaCl, 10 mM MgCl_2_, 2 mM TCEP, 1 *μ*M AMP-PNP, and protease inhibitor cocktail from Thermofischer Scientific). The lysate supernatant was bound to Ni-NTA agarose beads, washed with 25 mM imidazole Ni binding buffer, and eluted with 250 mM imidazole Ni elution buffer. Relevant fractions were pooled and diluted with Ni binding buffer adjusted to pH 8.0, such that the imidazole concentration was below 200 mM, for improved binding to MagStrep “type3” XT agarose beads (IBA lifesciences). Ni elutions were passed through XT beads at a low flow rate (∼1 ml/min), washed with a small volume of Ni binding buffer, then eluted using Strep XT elution buffer (100 mM HEPES, 150 mM NaCl, 10 mM MgCl_2_, 2 mM TCEP, 50 mM D+ biotin, pH 8.0). The concentration of protein in this elution was determined via Bradford assay, then ATP was added to a final concentration of 20 mM. This volume was supplemented with birA in a 50:1 MEK:birA ratio, then was incubated at 4 °C overnight. Following biotinylation overnight, the elution volume was concentrated to 5 ml using an Amicon Ultra concentrator (30 MWCO, Millipore) then further purified via size exclusion chromatography on a Superdex 200 Increase 10/300 column. Substrate quality was evaluated by mass spectroscopy and comparison of TR-FRET activity profiles to in-house standards.

### CRAF Expression and Purification for Crystallization

For crystallographic studies, His_6_-MBP-TEV-CRAF^337-615,Y340/341D^ and MEK1^SASA^ were co-expressed and purified as described above. Cells were lysed using sonication in buffer A (50 mM Tris-HCl at pH 7.4, 300 mM NaCl, 0.5 mM TCEP, 2 mM MgCl_2_, and 50 *μ*M ATP) supplemented with 25 mM imidazole and a protease inhibitor cocktail (Thermo Fisher Scientific). Lysates were clarified by ultracentrifugation and loaded onto a HisTrap HP column (Cytiva). After washing with buffer A supplemented with 40 mM imidazole, bound proteins were eluted using a linear gradient of buffer B (50 mM Tris-HCl pH 7.4, 300 mM NaCl, 0.5 mM TCEP, 2 mM MgCl_2_, 50 *μ*M ATP, and 500 mM Imidazole). Eluted fractions were incubated with TEV protease at a 1:50 molar ratio overnight at 4 ℃. Following protease digestion, proteins were buffer-exchanged to buffer C (50 mM Tris-HCl pH 7.0, 50 mM NaCl, 0.5 mM TCEP, 2 mM MgCl2, and 50 *μ*M ATP) using a HiPrep Desalting column packed with Sephadex G-25 resin (Cytiva) and quickly applied to a HiTrap SP HP cation exchange chromatography column (Cytiva). Cleaved His-MBP and excess MEK1 were removed in the flowthrough, and the remaining proteins were eluted using a linear gradient of buffer A. Eluted fractions were injected onto a HiLoad 16/600 Superdex 200 pg column (Cytiva) equilibrated with buffer D (50 mM HEPES-NaOH pH 7.5, 150 mM NaCl, and 1 mM TCEP). The purified proteins were concentrated to 20 mg/ml using an Amicon Ultra concentrator (30 kDa MWCO, Millipore).

### Crystallization and structure determination

Purified CRAF^337-615,Y340/341D^:MEK1^SASA^ proteins (20 mg/ml, 250 *μ*M) were incubated with 10 mM MgCl_2_, 1 mM AMPPNP, 1 mM PLX4720, and 1 mM CH5126766 at 4 ℃ overnight and crystallized at 22 ℃ using the sitting drop vapor diffusion method by mixing 150 nL proteins and 150 nL crystallization solutions. Initial crystals were obtained under commercial crystallization screening conditions containing 2% Tacsimate pH 5, 0.1 M Sodium Citrate pH 5.6, and 16% PEG 3350. Crystals were optimized in 1.4 ul hanging drops over wells containing 12−20 % PEG3350 and 2−6 % Tacsimate pH 5−6 with micro-seeds. The microseeds of crystals were prepared from the initial crystals using Seed Bead Kits (Hampton Research). Crystals were cryoprotected in Paratone oil (Hampton Research) and flash frozen in liquid nitrogen. X-ray diffraction data were collected at 100 K to a resolution of 2.91 Å on beamline 17-ID-1 (AMX) at the National Synchrotron Light Source II. Data were indexed and scaled using XDS (61). The structure was determined by molecular replacement with Phaser-MR (62) in the Phenix software suite (63), using the previously reported crystal structure of CRAF:MEK1 (PDB ID: 9AY7) as an initial search model. Phaser-MR identified four copies of CRAF:MEK1 complex in the asymmetric unit. The resulting model was further refined through iterative cycles of manual model building in Coot (64) and refinement using Phenix.Refine (65). Model quality was monitored using an R_free_ value calculated from 5% of reflections (66), and the final model was validated with MolProbity (67).

In order to increase the occupancy of AMPPNP in MEK1, purified CRAF^337-615,Y340/341D^:MEK1^SASA^ proteins (16 mg/ml, 200 *μ*M) were incubated with 10 mM MgCl_2_, 2 mM AMPPNP, 1 mM PLX4720, and 1 mM CH5126766, and crystallized as described above. X-ray diffraction data were collected to 3.50 Å at beamline 17-ID-1 (AMX), and structure determination followed the same procedure. The resulting electron density map showed full occupancy of AMPPNP in MEK1. The structures have been deposited in the Protein Data Bank with the accession codes 9O0U (CRAF:MEK1 with PLX4720 and CH5126766) and 9O0V (CRAF:MEK1 with PLX4720, CH5126766, and AMPPNP). The statistics of data collection and refinement are summarized in Supplementary Table 1. Protein structures are available in the Protein Data Bank under accession codes 9O0U and 9O0V.

## Data Availability

Protein structures have been deposited into the Protein Data Bank with the codes 9O0U and 9O0V, all other data are included in the manuscript and supplemental materials.

## Supporting Information

This article contains supporting information.

## Author Contributions

E. Tkacik performed enzyme inhibition assays. D. Jang and K. Boxer purified and crystallized proteins for structure determination. D. Jang determined crystal structures. E. Tkacik, B.H. Ha, J. Vinals, D. Jang and K. Boxer purified proteins for inhibitor characterization. M.J. Eck supervised the research. E. Tkacik, D. Jang and M.J. Eck wrote the manuscript. All authors reviewed and approved this manuscript prior to publication.

## Funding and Additional Information

This work was supported by NIH grants R35CA242461 (M.J.E.), P50CA165962 (M.J.E.), and by the PLGA fund of the Pediatric Brain Tumor Foundation. E.T. is the recipient of a graduate research fellowship from the Chleck Family Foundation. This work was conducted using resources at beamline 17-ID-1 of the National Synchrotron Light Source II, a U.S. Department of Energy (DOE) Office of Science User Facility operated for the DOE Office of Science by Brookhaven National Laboratory under Contract No. DE-SC0012704. The Center for BioMolecular Structure (CBMS) is primarily supported by the National Institutes of Health, National Institute of General Medical Sciences (NIGMS) through a Center Core P30 Grant (P30GM133893), and by the DOE Office of Biological and Environmental Research (KP1607011).

