## Supplemental Materials for "Characterization and inhibitor sensitivity of ARAF, BRAF, and CRAF complexes"

**Supplemental Table 1. Data collection and refinement statistics.***

|  | **CRAF/MEK1 with PLX4720 and CH5126766** | **CRAF/MEK1 with PLX4720, CH5126766, and AMPPNP** |
| --- | --- | --- |
| PDB Accession | 9O0U | 9O0V |
| **A. Data collection** |  |  |
| X-ray source | NSLS-II 17-ID-1 (AMX) | NSLS-II 17-ID-1 (AMX) |
| X-ray wavelength (Å) | 0.920201 | 0.920194 |
| Data process suite | *XDS* | *XDS* |
| Space group | *P*6_5_22 | *P*6_5_22 |
| Unit cell  a, b, c (Å)  α, β, 𝛾 (°) | 180.99, 180.99, 367.02  90.00, 90.00, 120.00 | 180.94, 180.94, 366.83  90.00, 90.00, 120.00 |
| Resolution range (Å) | 48.22 – 2.91  (2.97 - 2.91) | 48.20 – 3.50  (3.58 - 3.50) |
| Total reflections | 705,458 (47,055) | 300,934 (19,752) |
| Unique reflections | 147,295 (10,649) | 45,366 (3,155) |
| Multiplicity | 4.8 (4.4) | 6.6 (6.3) |
| Completeness (%) | 99.7 (97.4) | 99.5 (96.1) |
| Mean I/sigma(I) | 9.45 (1.43) | 12.57 (2.09) |
| *R_observed_* | 0.13 (1.19) | 0.14 (0.89) |
| *R_meas_* | 0.15 (1.35) | 0.15 (0.97) |
| CC1/2 (%) | 99.6 (43.5) | 99.7 (72.6) |
| **B. Model refinement** |  |  |
| *R_work_* / *R_free_*^a^ | 0.2302/0.2639 | 0.2054/0.2537 |
| No. of non-hydrogen atoms |  |  |
| protein | 18,489 | 18,507 |
| ligands | 271 | 365 |
| solvent | 44 | 8 |
| R.m.s. deviations |  |  |
| bond lengths (Å) | 0.007 | 0.004 |
| bond angles (°) | 1.086 | 0.825 |
| Ramachandran |  |  |
| favored (%) | 95.17 | 95.04 |
| outliers (%) | 0.04 | 0.13 |
| Rotamer outliers (%) | 0.69 | 0.00 |
| Clashscore | 11.18 | 8.5 |
| Average B-factor | 80.51 | 107.27 |
| proteins | 80.72 | 107.28 |
| ligands | 69.03 | 107.55 |
| water | 65.84 | 58.93 |

*Statistics for the highest-resolution shell are shown in parentheses.

^a^*R_free_* is calculated for a randomly chosen 5% of reflections that were not used for structure refinement.

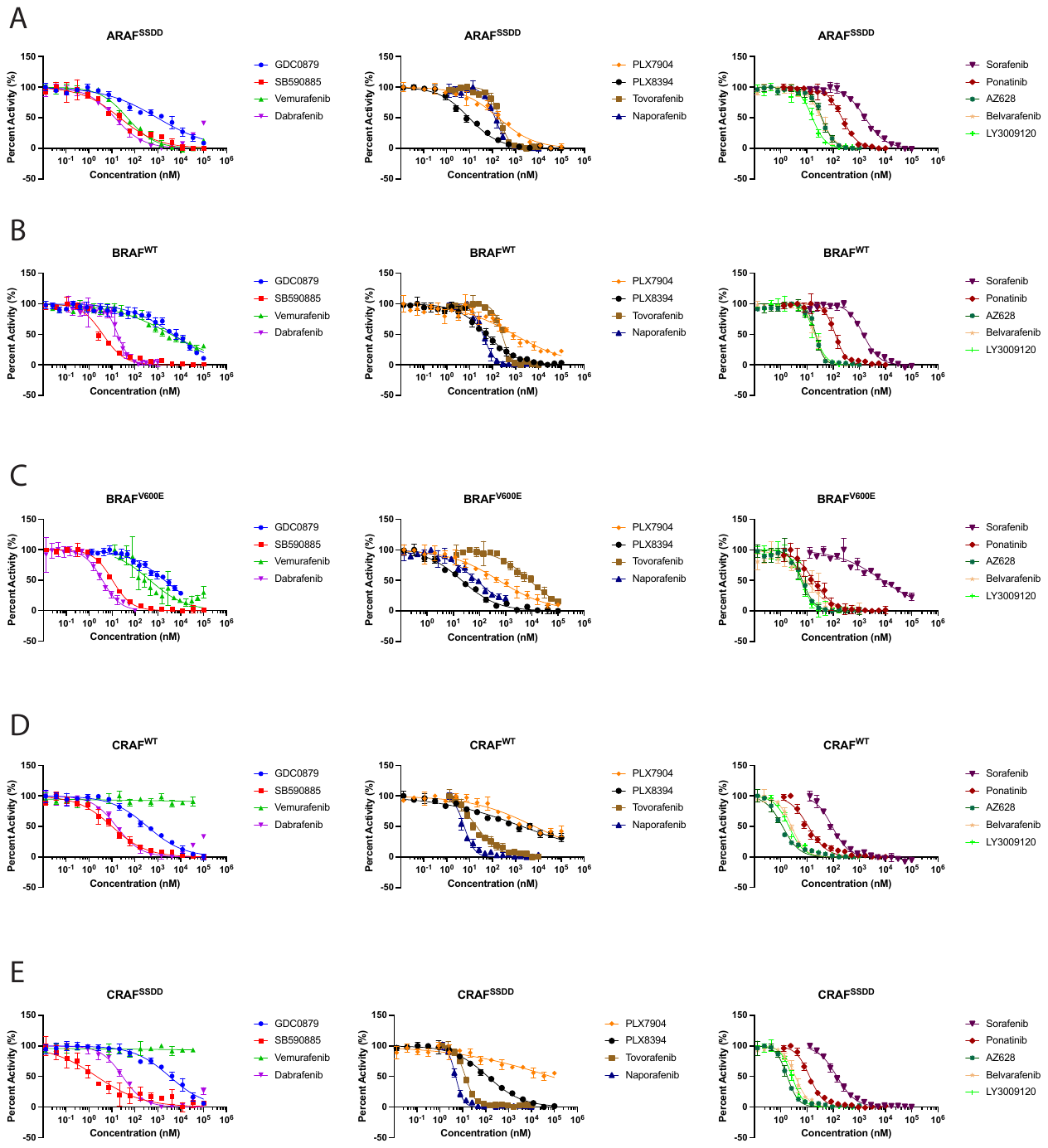

**Supplemental Figure 1. Concentration-response curves of RAF inhibitors studied here against ARAF^SSDD^, BRAF^WT^, BRAF^V600E^, CRAF^WT^, and CRAF^SSDD^.** Inhibition curves where the y-axis shows percent activity (%) for the indicated inhibitors for ARAF^SSDD^ (A), BRAF^WT^ (B), BRAF^V600E^ (C), CRAF^WT^ (D), and CRAF^SSDD^ (E). 100% activity corresponds to activity with no inhibitor while 0% activity is from assay background signal while fully inhibited. X-axis is inhibitor concentration, measured in nM. Data are plotted as mean ± SD from one independent experiment performed in triplicate (n = 3).

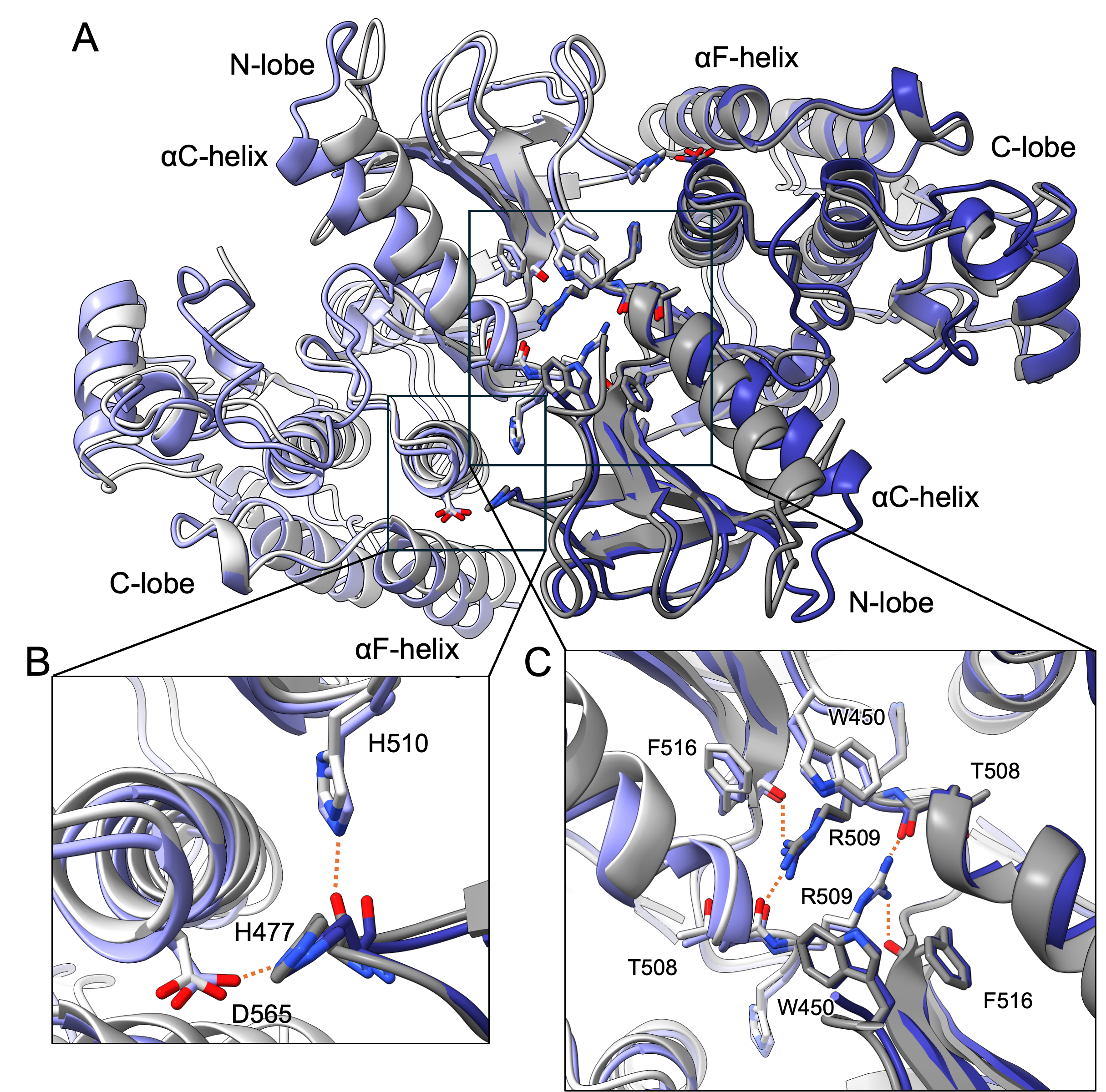

**Supplemental Figure 2. Prior BRAF-dabrafenib structure forms BRAF dimer with an unperturbed dimerization interface.** A, Comparison of inhibitor free BRAF dimer in blue (PDB ID: 4mne) and dabrafenib bound BRAF dimer in grey (PDB ID: 5csw). Kinase N-lobe, C-lobe are labelled. Helices with different conformations between the structures, αC and αF are also labelled. B and C, Zoomed in views of the boxed regions of the dimer interface, showing that despite differences in αC and αF conformations, the key dimer-dimer interface interactions are unperturbed in the dabrafenib bound structure. Key side chains are labelled and shown as sticks.

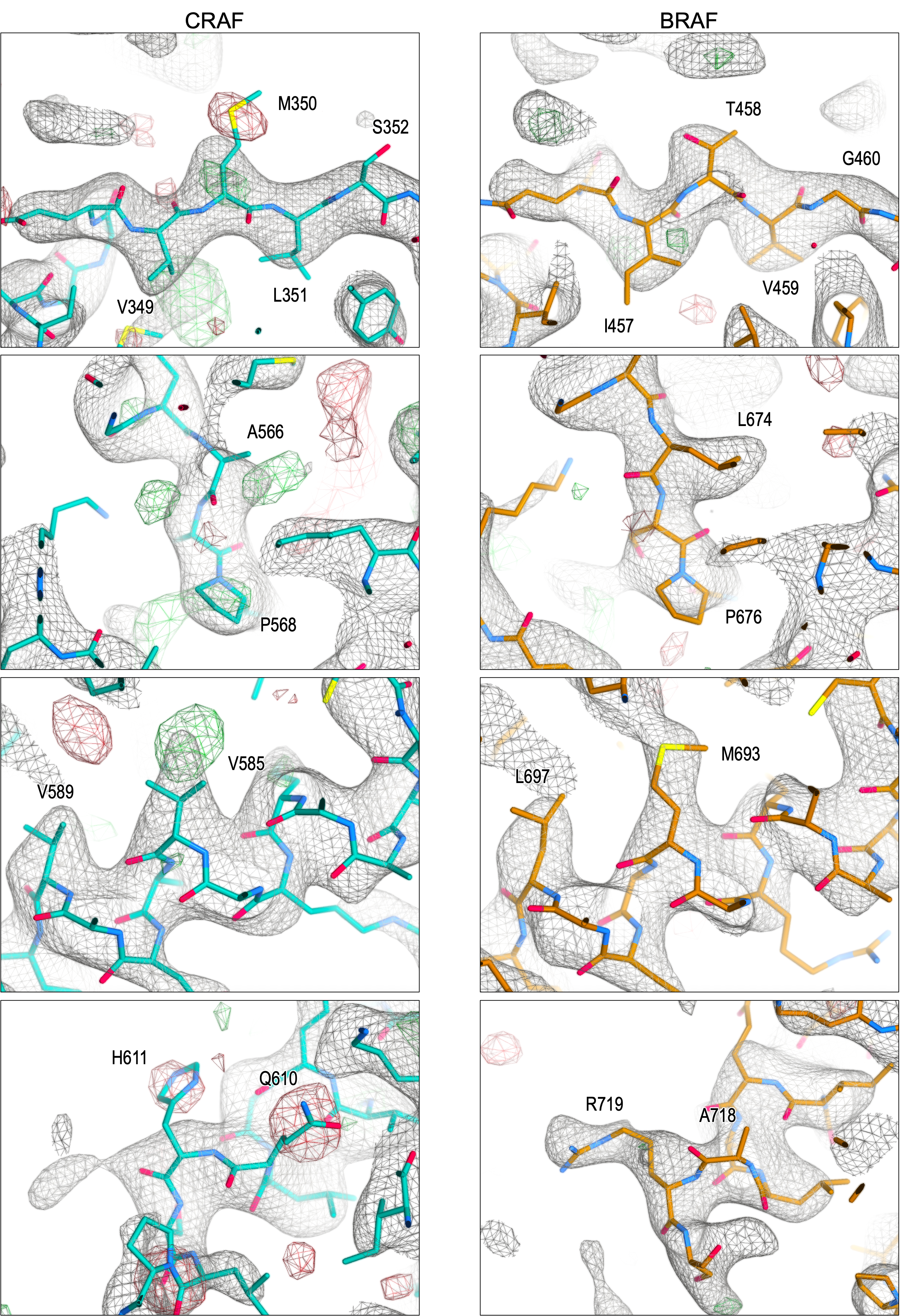

**Supplemental Figure 3. Re-examination of prior CRAF kinase domain structure (PDB ID: 3OMV) indicates that it is a misinterpreted BRAF kinase domain.** Electron density with the originally modelled CRAF residues (left panels) compared to the equivalent BRAF residues (right panels) reveals that the BRAF model fits better into the electron density maps. The 2Fo−Fc maps (grey) and Fo−Fc maps (green for positive and red for negative) are shown at contour levels of 1.0σ and 2.5σ, respectively. We also note that a subsequently deposited BRAF crystal structure (PDB Entry 4YHT) is isomorphous with this entry, providing further evidence that the underlying data were recorded from a crystal containing the BRAF kinase domain.

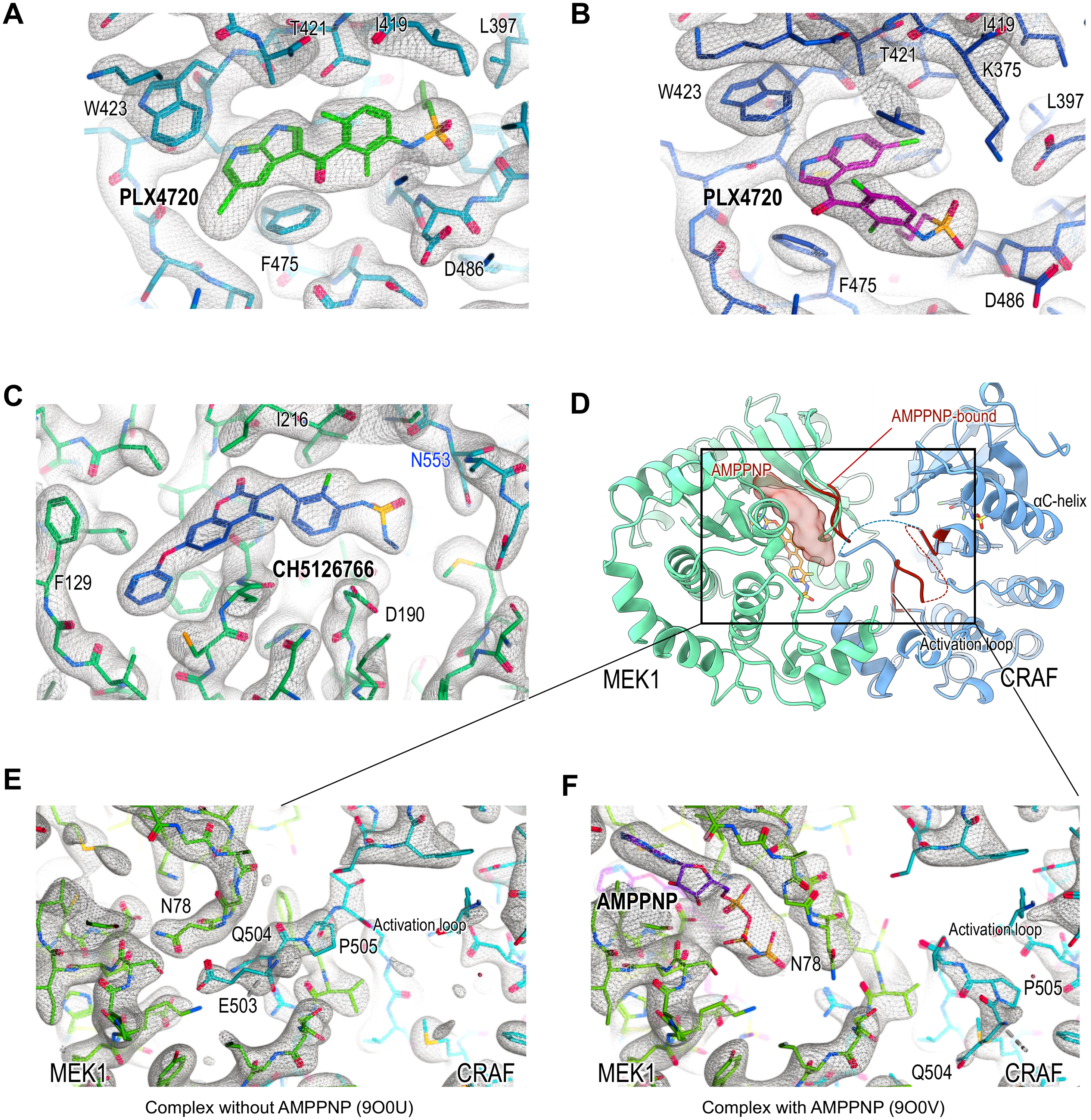

**Supplemental Figure 4. Electron density maps of inhibitors and nucleotide bound to the CRAF/MEK1 complex.** **A–C**, Crystal structure of CRAF/MEK1 with PLX4720 and CH5126766 (PDB ID: 9O0U). Canonical binding mode of PLX4720 (**A**) and non-canonical binding mode of PLX4720 (**B**) are shown in the ATP binding pocket of the CRAF kinase domain. The binding mode of CH5126766 (**C**) is shown in the allosteric pocket of MEK1. Maps shown in A–C are 2Fo−Fc maps (grey) at contour level of 1.6σ. **D**, Structural comparison of the CRAF:MEK1 complex with and without AMPPNP. The overall structure of CRAF:MEK1 without AMPPNP is shown, while AMPPNP and only the different regions of CRAF:MEK1 upon AMPPNP binding are shown as transparent surface and dark red cartoons, respectively. Dashed lines indicate the connecting segments in the CRAF activation loop. **E**–**F**,

Close-up views of the CRAF:MEK1 complex without AMPPNP (**E**) and with AMPPNP (**F**) overlayed with 2Fo−Fc maps at contour level of 1.3σ.
